# CSF sTREM2 correlates with CSF tau in advancing Parkinson’s disease

**DOI:** 10.1101/687269

**Authors:** Edward N. Wilson, Michelle S. Swarovski, Patricia Linortner, Marian Shahid, Abigail J. Zuckerman, Qian Wang, Divya Channappa, Paras S. Minhas, Siddhita D. Mhatre, Edward D. Plowey, Joseph F. Quinn, Cyrus P. Zabetian, Lu Tian, Frank M. Longo, Brenna Cholerton, Thomas J. Montine, Kathleen L. Poston, Katrin I. Andreasson

## Abstract

Parkinson’s disease (PD) is the second most common neurodegenerative disease after Alzheimer’s disease (AD) and affects 1% of the population above 60 years old. Although PD commonly manifests with motor symptoms, a majority of patients with PD subsequently develop cognitive impairment which often progresses to dementia, a major cause of morbidity and disability. PD is characterized by α-synuclein accumulation that frequently associates with amyloid beta (Aβ) and tau fibrils, the hallmarks of AD neuropathologic changes; this co-occurrence suggests that onset of cognitive decline in PD may be associated with appearance of pathologic Aβ and/or tau. Recent studies have highlighted the appearance of the soluble form of the Triggering Receptor Expressed on Myeloid cells 2 (sTREM2) receptor in CSF during development of AD. Given the known association of microglial activation with advancing PD, we investigated whether CSF and/or plasma sTREM2 increased with progression to PD dementia. We examined 165 participants consisting of 17 cognitively normal elderly, 45 PD patients with no cognitive impairment, 86 with mild cognitive impairment, and 17 with dementia. Stratification of subjects by CSF Aβ and tau levels revealed that CSF sTREM2 concentrations were elevated in PD subgroups with abnormal tau, but not Aβ, CSF concentration. These findings indicate that CSF sTREM2 could serve as a surrogate immune biomarker of neuronal injury in PD that is associated with cognitive decline.

**One sentence summary:** CSF sTREM2 correlates with CSF tau in PD

## Introduction

Parkinson’s disease (PD) affects 1% of the population above 60 years old (de Lau and Breteler, 2006) and is the most common neurodegenerative disease after Alzheimer’s disease (AD). While PD is typically considered a disease of motor function, one in four patients meet criteria for mild cognitive impairment (MCI) at diagnosis (Muslimovic *et al*., 2005; Aarsland *et al*., 2009) and nearly 80% eventually develop dementia during the course of the disease (Aarsland *et al*., 2003; Hely *et al*., 2008; Cholerton *et al*., 2013). These non-motor symptoms do not improve with dopamine enhancing therapies and contribute significantly to late morbidity, loss of quality of life, and mortality. Although at initial diagnosis, 6.5% of PD patients show abnormal CSF levels of Aβ_42_ and tau (Marek *et al*., 2018), at autopsy, 60-80% of PD subjects will have developed brain pathology consistent with AD, with prominent accumulation of amyloid plaques and/or tau containing neurofibrillary tangles (Tsuang *et al*., 2013; Dickson *et al*., 2018; Robinson *et al*., 2018). Thus in a vast majority of patients with PD, AD pathophysiologic processes are initiated at some point after diagnosis. Progression of parkinsonism and cognitive decline are accelerated in individuals with AD and/or cerebrovascular disease (CVD) (Tsuang *et al*., 2013; Dickson *et al*., 2018; Buchman *et al*., 2019). Identifying a biomarker that could help predict a change in cognitive function in PD would be a valuable tool for clinical management and outcome measure for clinical trials. Given the heterogeneous nature of PD, there is a significant unmet need for markers that could differentiate subgroups of PD patients that vary in rate of progression along cognitive and motor trajectories, or subgroups of PD patients with important differences in pathogenesis (Chen-Plotkin *et al*., 2018).

Triggering receptor expressed on myeloid cells 2 (TREM2) is an innate immune receptor expressed on the surface of monocytes, neutrophils, macrophages, and brain microglia (Colonna, 2003). Functionally, TREM2 plays a central role in phagocytosis of apoptotic neurons, misfolded proteins, and cellular debris (Neumann and Takahashi, 2007; Wang *et al*., 2016; Yeh *et al*., 2016). TREM2 signaling also enhances microglial survival, proliferation, chemotaxis, and through effects on metabolism, inhibits the microglial proinflammatory response (Hamerman *et al*., 2006; Turnbull *et al*., 2006; Otero *et al*., 2009; Ulland *et al*., 2017; Parhizkar *et al*., 2019). Overexpression of TREM2 attenuates neuroinflammation and protects dopaminergic neurons in the 1-methyl-4-phenyl-1,2,3,6-tetrahydropyridine (MPTP) mouse model of PD (Ren *et al*., 2018). In human genetic studies, the loss of function p.R47H variant of *TREM2* is a risk allele for AD (Guerreiro *et al*., 2013; Jonsson *et al*., 2013) and sporadic PD (Benitez *et al*., 2013; Rayaprolu *et al*., 2013).

The soluble form of TREM2 (sTREM2) is produced via the cleavage of membrane-bound TREM2 by disintegrin and metalloproteinase domain-containing protein (ADAM) family members, including ADAM10 and ADAM17 (Wunderlich *et al*., 2013; Kleinberger *et al*., 2014), and by alternative splicing (Guerreiro *et al*., 2013). CSF levels of sTREM2 are a sensitive marker of microglial activation and correlate with tau-mediated neuronal injury in AD (Kleinberger *et al*., 2014; Henjum *et al*., 2016; Heslegrave *et al*., 2016; Piccio *et al*., 2016; Suarez-Calvet *et al*., 2016a; Suarez-Calvet *et al*., 2016b; Suarez-Calvet *et al*., 2019). In this cross-sectional retrospective multicenter study, we tested whether CSF and/or blood sTREM2 appeared during the progression of PD to PD with dementia, and whether it correlated with CSF concentrations of Aβ_1-42_, Aβ_1-40_, total tau, and phospho-tau_p181_, or with cognitive status.

## Materials and methods

### Study population

Participants (N=165) were recruited from the Pacific Udall Center (PUC) cohort consisting of sites at Seattle (VA Puget Sound Health Care System/University of Washington) and Portland (Cholerton *et al*., 2013) and from the Stanford Movement Disorders Clinic. Participants were included if they were cognitively normal healthy adults or if they met United Kingdom Parkinson’s Disease Society Brain Bank clinical diagnostic criteria for PD, had a cognitive diagnosis assigned (no cognitive impairment, MCI (Cholerton *et al*.), PD dementia (Cholerton *et al*.), and if they completed lumbar puncture. A full neuropsychological battery assessing multiple domains was given to assign a cognitive diagnosis. Participants from Stanford were defined as cognitively impaired if scores were ≥1.5 standard deviations below age- and education-matched normative values on at least two separate neuropsychological measures, regardless of domain (Hendershott *et al*., 2019). The Pacific Udall Center assigned cognitive diagnoses at a clinical consensus conference and required evidence of subjective and observed cognitive decline. For all PD participants, cognitive impairment was further classified as PD-Dementia, as opposed to PD-MCI if the impairment was severe enough to interfere with daily activities (Emre *et al*., 2008). All study protocols were approved by Institutional Review Boards of Stanford University, Oregon Health & Science University, or VA Puget Sound Health Care System/University of Washington. In accordance with the Declaration of Helsinki, written informed consent was obtained from each study participant or their legally authorized representative.

### Global cognitive function

Global cognitive function was assessed using the Montreal Cognitive Assessment (MoCA) test (Nasreddine *et al*., 2005). The MoCA is the most commonly used cognitive assessment in PD and has a high sensitivity and specificity for identifying MCI in PD (Hendershott *et al*., 2017). The MoCA is used in large multicenter PD studies because it assesses primary cognitive domains at risk in PD, has been validated in multiple languages, and requires substantially less time and training to properly administer and score than a full neuropsychological assessment (Chou *et al*., 2010).

### Plasma and CSF collection

For the Stanford University cohort, fasted plasma was collected within two weeks of lumbar puncture. For the two PUC cohorts, centers in Seattle and Portland performed CSF and plasma blood draws on the same morning. CSF samples were subjected to a maximum of two freeze-thaw cycles, as recommended by Consensus of the Task Force on Biological Markers in Psychiatry of the World Federation of Societies of Biological Psychiatry (Lewczuk *et al*., 2018).

### Measurement of AD CSF core biomarkers

The following cutoff values for abnormal CSF biomarkers were employed: ratio CSF Aβ_1-42_/Aβ_1-40_ < 0.10, phospho-tau_181_ > 40 pg/ml, and total tau > 456 pg/ml, as measured using Lumipulse G Assays (Fujirebio, Malvern, PA) on the LUMIPULSE G fully automated platform (Bayart *et al*., 2019; Paciotti *et al*., 2019). CSF samples from all three cohorts were measured all together in one day by trained operators who were blinded to the clinical information.

### CSF and plasma sTREM2 measurement

Electrochemiluminescent immunoassays (ECLIAs) on the Meso Scale Discovery platform (Meso Scale Discovery, Rockville, MD) were used to measure CSF and plasma sTREM2. MSD GOLD 96-Well Streptavidin plates were blocked with 3% BSA and coated with a solution containing 0.25 µg/ml biotinylated polyclonal goat anti-human TREM2 capture antibody (BAF1828, R&D Systems, Minneapolis, MN). CSF and plasma samples diluted 1:4 in 1% BSA were added to the prepared plates and incubated overnight at 4°C. Monoclonal mouse anti-human TREM2 detection antibody (B-3, sc373828, Santa Cruz Biotechnology, Santa Cruz, CA) was added at a concentration of 1 µg/ml, followed by SULFO-TAG anti-mouse secondary antibody (1 µg/ml, R32AC-5, MSD). Plates were washed 4 times between incubations with PBS/0.05% Tween-20 (PBS-T) buffer. Samples were distributed in a randomized manner across plates and read in duplicate by an operator blinded to clinical information. Plates were read using a MESO QuickPlex SQ 120 running Discovery Workbench v4 software (MSD). CSF and plasma sTREM2 concentrations were calculated using the standard curve generated for each plate using recombinant human TREM2 protein (Sino Biological, Wayne, PA). A dedicated CSF and plasma sample was loaded onto all plates and used to normalize values. Interplate CVs were <15%. Samples were measured on the same day using the same reagents.

### Statistical analysis

CSF (Shapiro-Wilk normality test, W = 0.8499, *P* < 0.0001) and plasma sTREM2 (W = 0.2202, *P* < 0.0001) did not follow a normal distribution and were log_10_-transformed to approach the assumptions of Gaussian normal distribution. Therefore, all statistical analyses described in this study are performed with the log_10_-transformed values. Categorical variables were assessed by performing the chi-square test. Association of sTREM2 with continuous variables was evaluated using a linear regression model. Demographic parameters between groups were evaluated using one-way ANOVA followed by Tukey corrected *post hoc* pairwise comparisons (for parametric data) and Mann-Whitney U test or Kruskal-Wallis test followed by Dunn’s corrected *post hoc* comparisons (for non-parametric data). To determine whether CSF and plasma sTREM2 differed among clinically- and biomarker-defined groups, Log_10_-transformed CSF sTREM2 levels were analyzed using ANCOVA with clinical diagnosis as fixed factor and age and gender as covariates. Tukey’s multiple comparisons test was performed for *post hoc* testing. Receiver operating characteristic (ROC) curve analysis was performed to assess CSF sTREM2 in differentiating PD participants with abnormal CSF tau concentration from those with normal CSF tau concentration. Association between CSF sTREM2 and core AD CSF biomarker was assessed using linear mixed-effects model adjusting for age and gender as fixed effects and CSF sample freeze-thaw number as random effect. Association between CSF sTREM2 and MoCA score was assessed using a linear model adjusted for age, gender, years of education, and study site. Statistical tests were performed using GraphPad Prism software (GraphPad Inc, La Jolla, CA) and the freely available statistical software R (http://www.r-project.org/). All tests were two-sided and a significance level of α = 0.05 was adopted.

### Data availability

The data that support the findings of this study are available from the corresponding author, upon request.

## Results

### Study participant characteristics

Demographic information for the study population is presented in **Table 1**. The clinically-defined diagnostic groups consisted of healthy cognitively unimpaired age-matched controls (Healthy Control), cognitively normal PD subjects (PD-Normal), PD subjects with mild cognitive impairment (PD-MCI) and PD subjects with dementia (PD-D). These groups differed in gender distribution (χ^2^ = 16.44, df = 1, *P* < 0.0001), with a larger proportion of male subjects in the PD-MCI and PD-D groups. This is in line with previous data indicating that PD patients who are male are more likely to be cognitively impaired (Cholerton *et al*., 2018). No differences were observed between any of the diagnostic groups for age (one-way ANOVA: *F*_(3,161)_ = 2.526; *P* > 0.05) or disease duration as defined as time in years since diagnosis (Kruskal-Wallis χ^2^ = 4.767, df = 3, *P* = 0.092). As expected, performance on the MoCA, a test sensitive to global cognition and executive function in PD (Hendershott *et al*., 2017), differed between clinical diagnostic groups (Kruskal-Wallis chi-squared = 67.522, df = 3, *P* < 0.0001). While no significant difference was observed between Healthy Control and PD-Normal, the MoCA score was significantly lower in the PD-MCI (*P* < 0.001) and PD-D (*P* < 0.0001) groups. We observed significant differences in the off-medicine Movement Disorders Society-Unified PD Rating Scale motor part III (MDS-UPDRS) across PD diagnoses (Kruskal-Wallis test (χ^2^= 15.69, df=3, *P* = 0.0004), which confirmed equal motor symptom severity in the PD-Normal and PD-MCI groups *(P* > 0.05), but increased motor symptoms in the PD-D compared to both PD-Normal (*P* < 0.01) and PD-MCI (*P* < 0.001) groups. Finally, the percentage of *APOE* ε4 carriers was not significantly different in any of the diagnostic groups (χ^2^ = 0.419, *P* = 0.518).

**Table 1.** Data Summary Table

Figures and Legends
| Table 1. Data Summary Table |  |  |  |  |  |  |
| --- | --- | --- | --- | --- | --- | --- |
| Variable | Total (n = 165) | Healthy Control (n=17) | PD-Normal (n=45) | PD MCI (n=86) | PD Dementia (n=17) | P-value |
| Gender (F/M), no. (%) | 52 (32)/113 (68) | 10 (59)/7 (41) | 22 (49)/23 (51) | 17 (20)/69 (80) | 3 (18)/14 (82) | P<0.0001, a |
| Age, years (Mean, SD) | 66.8 (8.32) | 65.06 (6.7) | 64.4 (7.6) | 67.8 (8.3) | 69.5 (10.3) | ns,b |
| Disease duration, years (Mean, SD) |  |  | 4.6 (3.9) | 5.6 (5.6) | 7.5 (4.8) | ns,c |
| MoCA Score (Mean, SD) | 24.6 (3.9) | 27.4 (2.0) | 27.1 (2.3) | 24.1 (2.5)*.^ | 17.8 (5.1)#,^,@ | P<0.0001,c |
| MDS-UPDRS, part III Off (Mean, SD) | 26.8 (16.4) | 1.18 (1.5) | 29.1 (14.6) | 27.5 (13.8) | 43.2 (12.5)\$,@ | P<0.0001, c,d |
| APOE e4 allele (% Carriage) | 23.0 | 17.6 | 32.6 | 19.8 | 20.0 | ns, a |

a Chi-Square test; b One Way ANOVA Post Hoc significances for age: PD-Normal versus PD Dementia P < 0.05; c Kruskal-Wallis test; d Kruskal-Wallis test comparing PD subgroups only; \*, P<0.001 vs Control; #, P<0.0001 vs Control; ^, P<0.0001 vs PD-Normal; \$, P<0.01 vs PD-Normal; &, P<0.01 vs PD-MCI; @, P<0.001 vs PD-MCI.

### Association between sTREM2 with demographic information

Overall, age was positively associated with CSF sTREM2 (β = +0.336, SE = 0.006, *P* < 0.0001) but not plasma sTREM2 (β = −0.064, SE = 0.016, *P* = 0.412). We observed no association between disease duration and CSF sTREM2 (β = +0.029, SE = 0.010, *P* = 0.714) or plasma sTREM2 (β = +0.0166, SE = 0.027, *P* = 0.534). Furthermore, compared to female participants, male participants showed significantly higher CSF sTREM2 (Mann-Whitney U = 1660, N=165, *P* < 0.0001) but not plasma sTREM2 (Mann-Whitney U = 2783, N = 165, *P* > 0.05). For these reasons, we included age and gender in our statistical models for CSF sTREM2. Including age and gender in statistical models evaluating plasma sTREM2 did not change the outcome of the analyses.

### Baseline CSF and blood plasma sTREM2 levels in PD

We first determined whether CSF and plasma levels of sTREM2 differed in PD along the cognitive spectrum between PD-Normal, PD-MCI, and PD-D, compared to healthy age-matched controls. Log_10_-transformed CSF sTREM2 levels were analyzed using ANCOVA with clinical and cognitive diagnosis as fixed factor and age and gender, and study site as covariates. This revealed that CSF sTREM2 levels were significantly modulated in PD (One-way ANCOVA: *F*_(3,159)_ = 4.588; *P* = 0.004; Fig. 1A) with CSF sTREM2 significantly elevated in cognitively normal PD participants compared to healthy control subjects (Tukey’s multiple comparisons test: *P* = 0.046). Higher concentrations of sTREM2 were observed in plasma compared to CSF overall, however, no differences were observed between PD-Normal, PD-MCI, and PD-D (Fig. 1B) (One-way ANCOVA: *F*_(3,159)_ = 0.518; *P* = 0.671). Therefore, subsequent analyses focused on CSF sTREM2.

**Figure 1:**
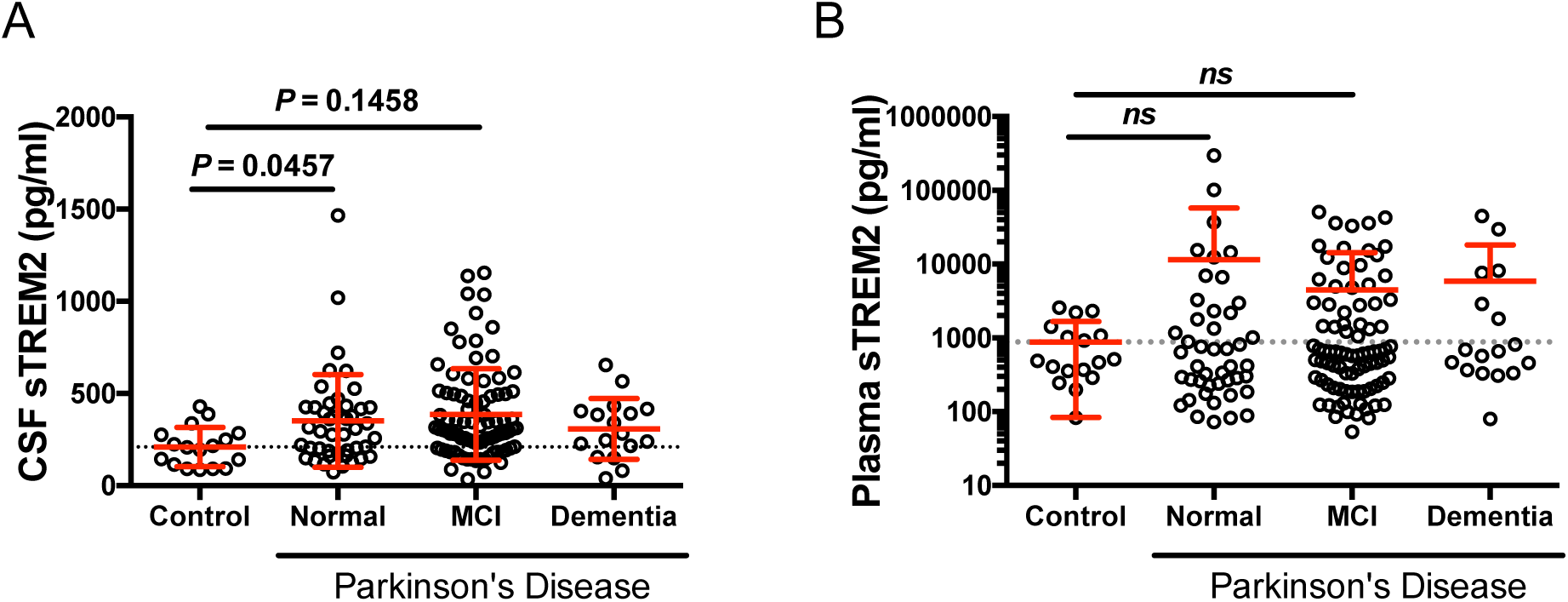
CSF and Plasma sTREM2 in healthy control subjects and in PD subjects separated based on cognitive status. (**A**) Scatter plot showing levels of CSF sTREM2 are elevated in PD Normal relative to Control. (**B**) Plasma sTREM2 concentration is not significantly different across any of the diagnostic groups. Solid bars represent the mean and standard deviation (SD). CSF and plasma sTREM2 data were log-transformed and analyzed using a one-way ANCOVA with age and gender as co-variates, followed by *post hoc* multiple comparisons using Tukey contrasts.

### Heterogeneity in amyloid and tau co-pathologies in Parkinson’s disease

CSF Aβ and tau concentrations can reveal biomarker-defined AD (Jack *et al*., 2018), even within clinically-defined PD patient populations (van Steenoven *et al*., 2016). Therefore, we used measurement of CSF amyloid and tau concentrations to stratify participants into one of five CSF biomarker-defined groups: 1) Controls with normal amyloid and tau (Controls); 2) PD with normal amyloid and tau [pure Lewy Body (LB) PD]; 3) PD with abnormal amyloid and normal tau (PD:Aβ+); 4) PD with both abnormal amyloid and tau (PD:Aβ+/Tau+); and 5) PD with normal amyloid but abnormal tau (PD:Tau+). Levels of CSF amyloid and tau biomarkers across each of the CSF biomarker-defined diagnostic groups are shown in **Table 2**.

**Table 2.** CSF Amyloid and Tau Biomarkers

|  | Controls (n = 17) | pLB PD (n=86) | PD: Aβ+ (n=16) | PD: Tau+ (n=23) | PD: Aβ+/Tau+ (n=23) | P-value* |
| --- | --- | --- | --- | --- | --- | --- |
| CSF Aβ1-42 (pg/ml; Mean, SD) | 1002.77 (390.94) | 1000.14 (334.19) | 596.75 (201.42) | 1748.83 (455.96) | 756.13 (322.66) | <0.0001 |
| CSF Aβ1-40 (pg/ml; Mean, SD) | 7987.59 (2853.18) | 7568.00 (2218.13) | 6981.81 (1974.63) | 12902.22 (2522.55) | 10146.17 (3107.54) | <0.0001 |
| CSF t-tau (pg/ml; Mean, SD) | 225.94 (90.70) | 217.12 (91.54) | 161.44 (67.37) | 382.13 (112.74) | 559.48 (303.61) | <0.0001 |
| CSF p-tau 181 (pg/ml; Mean, SD) | 33.37 (11.11) | 27.38 (6.52) | 38.01 (7.92) | 48.58 (8.53) | 67.81 (32.44) | <0.0001 |

\* Kruskal-Wallis test

### CSF sTREM2 is elevated in PD with abnormal CSF tau concentration

To determine the utility of a biomarker-defined diagnosis in PD, we next compared CSF sTREM2 expression across biomarker-defined groups (Fig. 2A). After adjusting for age and gender, we found that CSF sTREM2 was significantly different between the biomarker-defined groups (F_4,158_ = 8.926, *P* < 0.0001). *Post-hoc* testing revealed no significant differences between healthy control, pure LB PD, or PD:Aβ+ groups. In contrast, CSF sTREM2 was significantly elevated in the PD:Aβ+/Tau+ group compared to control (*P* < 0.001), pure LB PD (*P* < 0.001), and PD Aβ+ groups (*P* = 0.011). CSF sTREM2 remained significantly elevated in the PD:Tau+ group, even in the absence of abnormal CSF Aβ, compared to the control (*P* < 0.001), pure LB PD (*P* = 0.002), and trended towards elevated relative to the PD:Aβ+ (*P* = 0.057) groups. Receiver operating characteristic (ROC) curve analysis revealed that CSF sTREM2 could differentiate PD participants with abnormal CSF tau concentration (*N*=46) from those with normal CSF tau concentration (*N*=119; Fig. 2B), with an area under curve (AUC) of 0.828 (95% confidence interval (CI) = 0.757-0.899).

**Figure 2:**
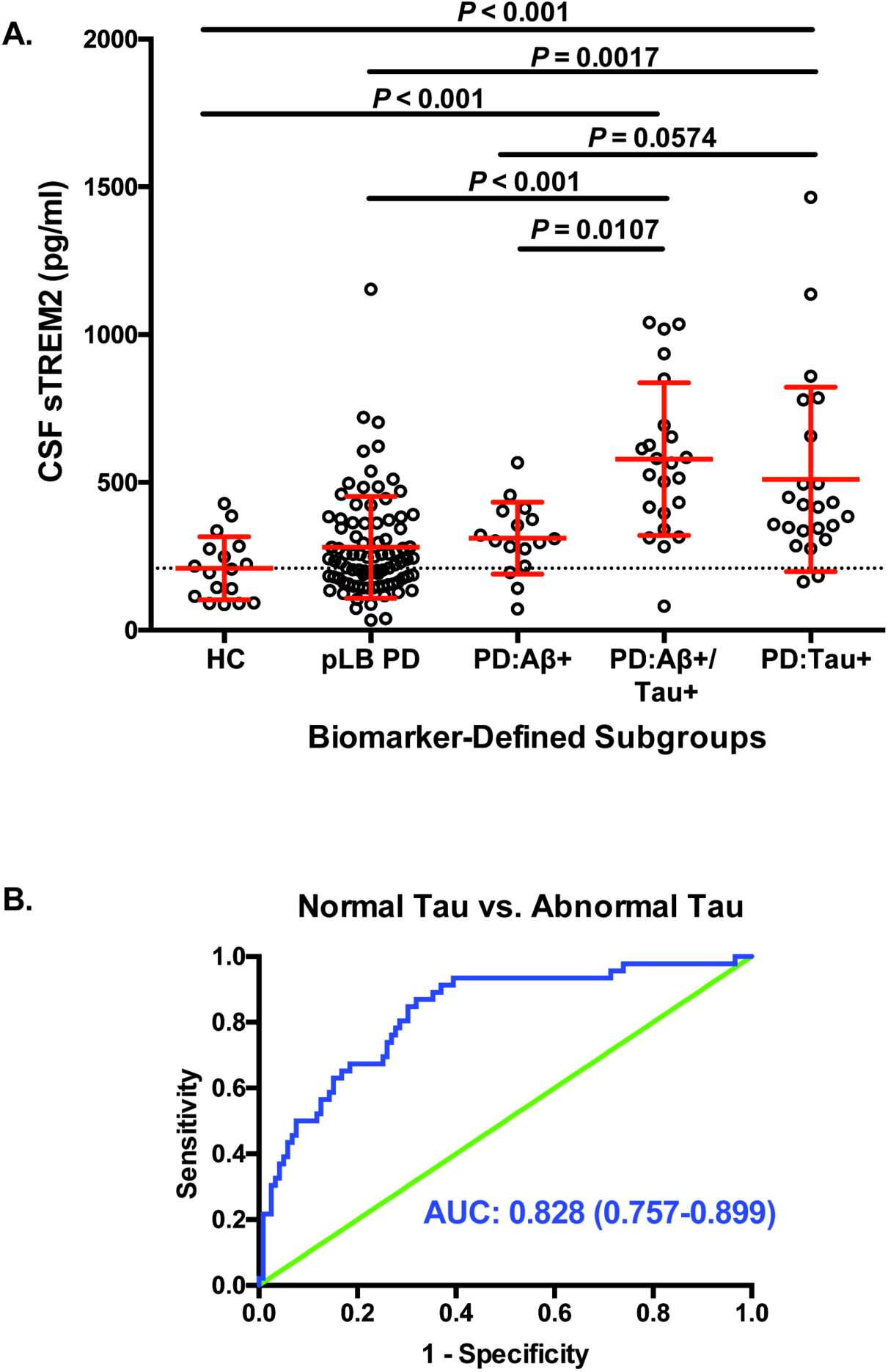
CSF sTREM2 concentration across biomarker-defined diagnostic profiles. **A**) CSF amyloid and tau concentrations were used to stratify participants into one of five CSF biomarker-defined groups: 1) Controls with normal amyloid and tau (HC); 2) PD with normal amyloid and tau [pure Lewy Body (LB) PD]; 3) PD with abnormal amyloid and normal tau (PD:Aβ+); 4) PD with both abnormal amyloid and tau (PD:Aβ+/Tau+); and 5) PD with normal amyloid but abnormal tau (PD:Tau+). Log-transformed CSF sTREM2 data were analyzed using a one-way ANCOVA adjusted by age and gender followed by Tukey contrasts *post hoc* multiple comparisons. Red bars indicate mean and SD. **B**) Receiver operating characteristic (ROC) curve analysis of CSF sTREM2 in discriminating PD with tau co-pathology from PD without tau co-pathology. The area under the curve was 0.828 (95% CI = 0.757–0.899).

### CSF sTREM2 in biomarker-defined PD groups across clinical diagnosis

CSF sTREM2 concentration was determined for each PD patient subgroup stratified according to CSF biomarker profile. After adjusting for age and gender, CSF sTREM2 concentrations were significantly different between biomarker-defined groups within the diagnoses PD-Normal (F_3,39_ = 4.415, *P* = 0.009), PD-MCI (F_3,80_ = 6.626, *P* = 0.001, and PD-D (F_2,12_ = 4.498, *P* = 0.035; (Fig. 3). *Post-hoc* analysis using Tukey contrasts revealed that CSF sTREM2 level was significantly higher in the PD:Aβ+/Tau+ and PD:Tau+ subgroups compared to the pure LB PD subgroup in all three PD-Normal, PD-MCI, and PD-D diagnoses. This indicates that CSF sTREM2 is elevated in PD patient subgroups with abnormal CSF tau concentration, but not CSF Aβ concentration, before the onset of MCI or dementia.

**Figure 3:**
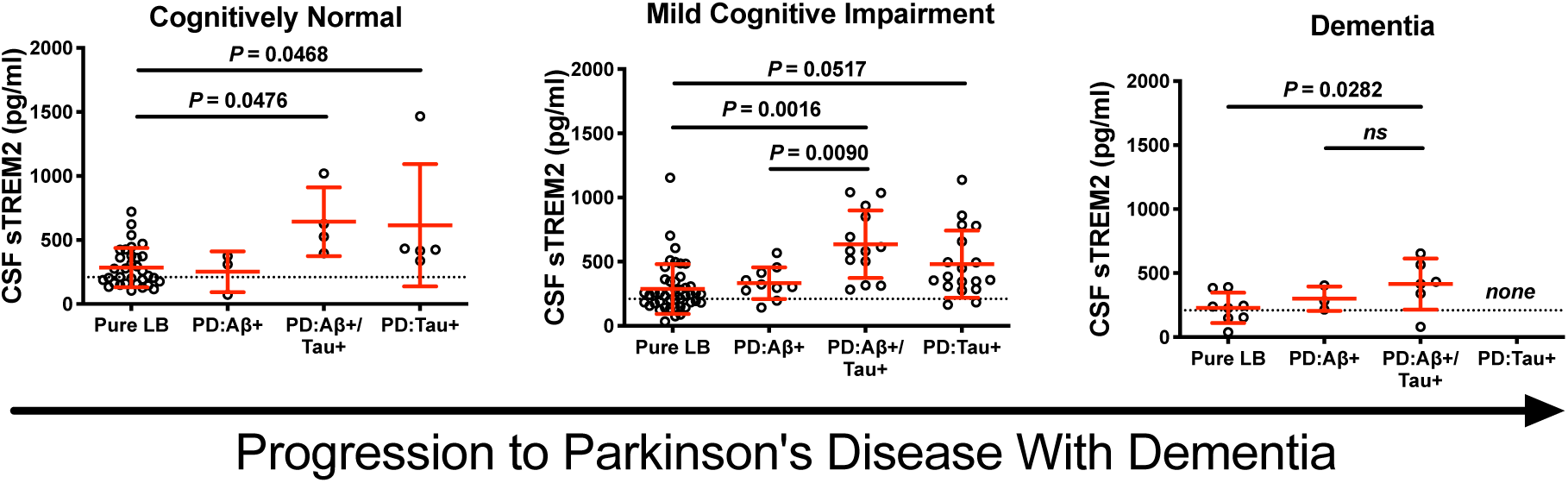
CSF sTREM2 according to abnormal Aβ and tau biomarker profile subgroups within each of the clinically-defined PD diagnoses. Scatter plot representing levels of CSF sTREM2 (log-transformed) in participants for each of the four biomarker subgroups defined by abnormal Aβ and/or tau within corresponding cognitive diagnosis. The PD-Dementia continuum spans normal (no cognitive impairment), through mild cognitive impairment, to PD with dementia. Log-transformed CSF sTREM2 data were analyzed using a one-way ANCOVA and were adjusted by gender and age followed by *post hoc* multiple comparisons using Tukey contrasts. Red bars indicate mean and SD.

### Association between CSF sTREM2 and CSF A**β** and tau concentrations

We next studied the relationship between CSF sTREM2 and the core AD CSF biomarkers phospho-tau_p181_, total tau, Aβ_1_-_42_, and Aβ_1_-_40_. We found that in the pooled group of participants, CSF sTREM2 concentration was positively associated with CSF total tau concentration in PD-Normal (β = +0.592, SE = 0.186, *P* = 0.003), PD-MCI (β = +0.510, SE = 0.118, *P* < 0.0001), and despite relatively few cases, tended towards association in the PD-D diagnosis (β = +0.511, SE = 0.267, *P* = 0.078); Fig. 4A-C). Likewise, positive associations between CSF sTREM2 and CSF phospho-tau_181_ were detected in the PD-Normal (β = +1.068, SE = 0.262, *P* = 0.0002), PD-MCI (β = +0.3490, SE = 0.156, *P* = 0.028), and PD-D groups (β = +0.477, SE = 0.177, *P* = 0.018; Fig. 4D-F). No association was observed between CSF sTREM2 and CSF Aβ_1-42_ in the PD-Normal (β = +0.327, SE = 0.209, *P* = 0.126), PD-MCI (β = +0.140, SE = 0.140, *P* = 0.320), or PD-D diagnosis (β = +0.214, SE = 0.318, *P* = 0.512); Fig. 4G-I). On the other hand, CSF sTREM2 was strongly associated with CSF Aβ_1-40_ in PD-Normal (β = +0.910, SE = 0.234, *P* = 0.0004) and PD-D groups (β = +0.797, SE = 243, *P* = 0.006), and tended towards a positive association in the PD-MCI (β = +0.329< SE = 0.177, *P* = 0.067; Fig. 4J-L).

**Figure 4:**
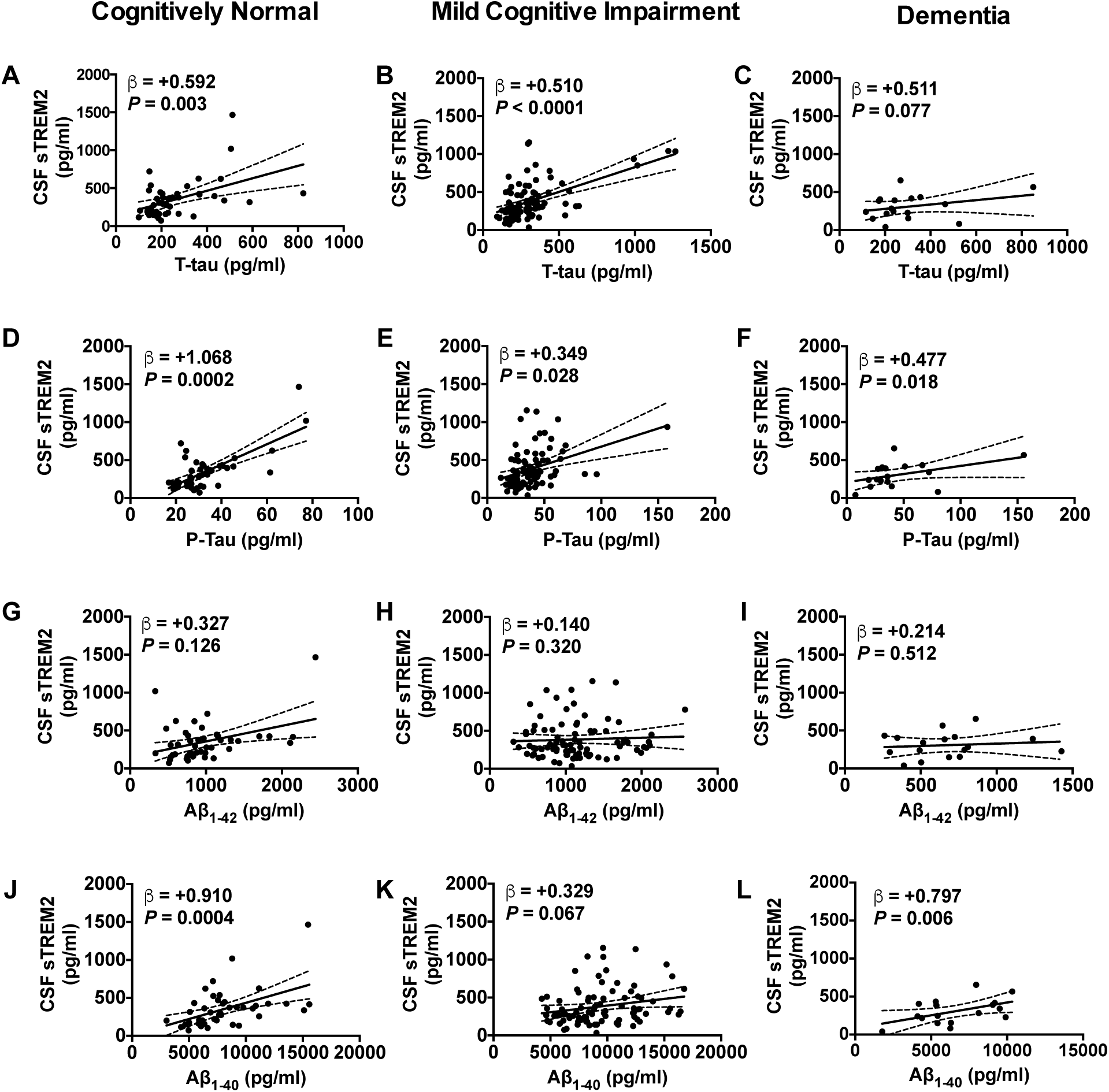
Association between CSF sTREM2 and CSF Aβ and tau biomarkers. Significant association was observed between CSF sTREM2 levels and CSF total tau (**A-C**) and p-Tau181 (**D-F**) for all diagnostic groups. In no group did CSF sTREM2 associate with Aβ_1-42_ (**G-I**). Significant association was observed between CSF sTREM2 levels and CSF Aβ_1-40_ for PD Cognitively Normal (**J**) and PD Dementia diagnoses (**L**), and trended towards significance in PD MCI (**K**). Associations between CSF sTREM2 and biomarkers were assessed using linear mixed-effects model with age and gender as fixed effects and freeze-thaw cycle number as random effect. Biomarker values were log-transformed to reduce skewness. Plotted is the 95% confidence band of the best-fit line from the linear regression. β estimates and *P*-values from the linear model are shown.

### CSF sTREM2 concentration is positively associated with MoCA in PD with abnormal CSF tau concentration

Finally, we evaluated whether CSF sTREM2 levels were associated with performance on the MoCA using a linear model adjusted for age, gender, years of education, and study site. No association between CSF sTREM2 and MoCA score was detected within biomarker-defined Pure LB PD (β = +0.079, SE = 0.012, *P* = 0.456) or PD:Aβ+ (β = +0.316, SE = 0.028, *P* = 0.348) subgroups (Fig. 5). In contrast, elevated CSF sTREM2 was associated with higher MoCA score in the PD:Aβ+/Tau+ (β = +0.545, SE = 0.028, *P* = 0.021) and the PD:Tau+ (β = +0.538, SE = 0.059, *P* = 0.036) subgroups.

**Figure 5:**
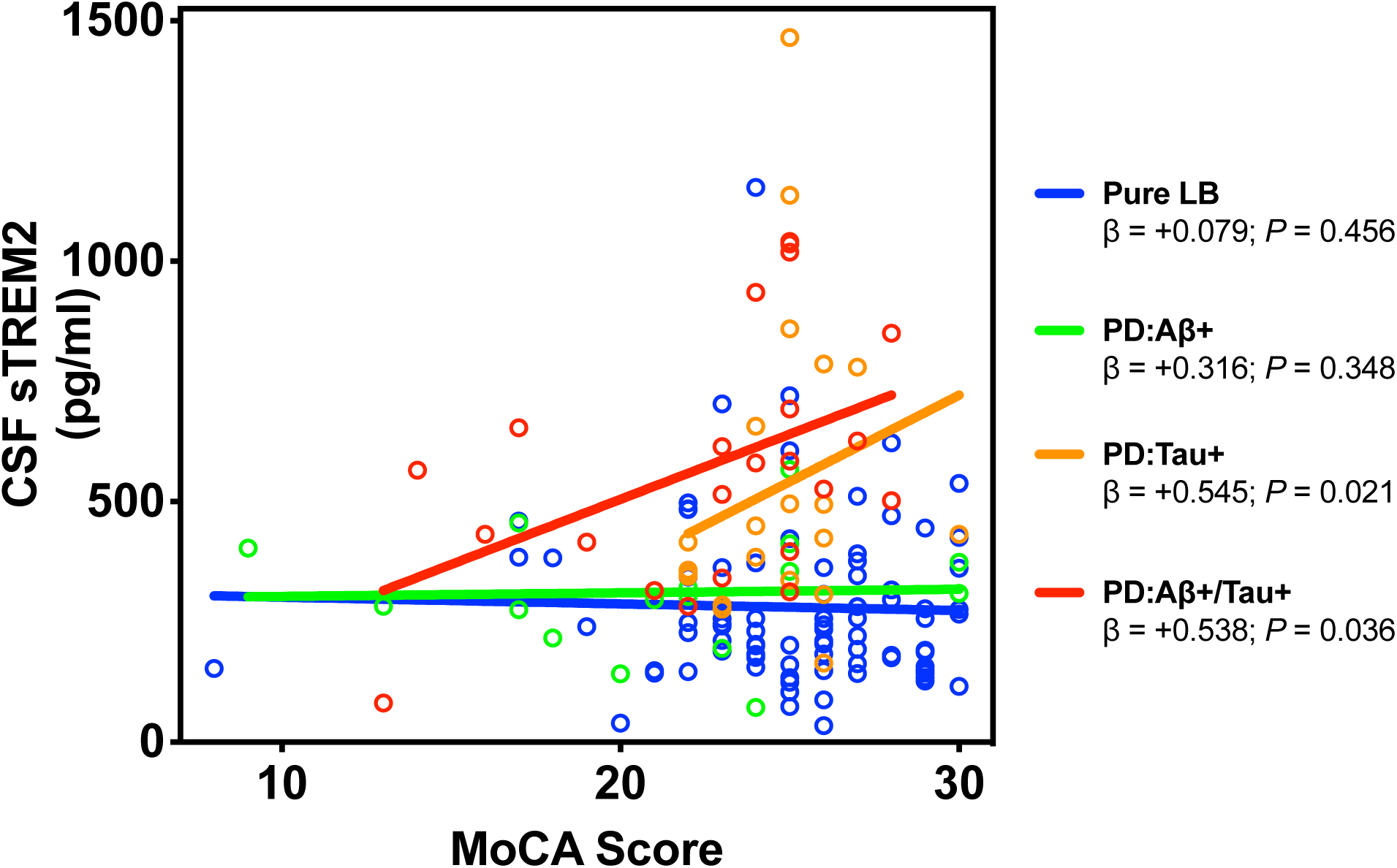
Association between CSF sTREM2 and MoCA score in the biomarker PD patient subgroups. A significant positive association between CSF sTREM2 (log-transformed) and MoCA score was detected in the PD:Tau+ and PD:Aβ+/Tau+ patient subgroups. Plotted is the linear regression for each group. β estimates and *P*-values from the linear model adjusted by age, gender, years of education, and study site are shown.

## Discussion

Substantial biological heterogeneity underlies the clinical presentation and progression of PD and there is a significant need for markers that differentiate subgroups of PD patients in their rate of progression along cognitive and motor trajectories, or in the types of co-morbid disease (Chen-Plotkin *et al*., 2018). We report here that CSF sTREM2 is increased in PD patient subgroups with abnormal CSF tau concentration, with or without abnormal CSF Aβ_1-42_ concentration, and that this increase precedes cognitive impairment.

Our observation that CSF sTREM2 is increased prior to cognitive symptoms in PD stages is similar to that seen in the progression to late-onset AD (Kleinberger *et al*., 2014; Henjum *et al*., 2016; Heslegrave *et al*., 2016; Piccio *et al*., 2016; Suarez-Calvet *et al*., 2016b; Suarez-Calvet *et al*., 2019), and early-onset familial AD (Suarez-Calvet *et al*., 2016a). In AD, the increase in CSF sTREM2 concentration occurs after pathological decrease in CSF Aβ_1-42_ levels, and coincident with increased CSF tau concentration. In PD, we observed a similarly strong association between CSF sTREM2 and phospho-tau_181_ and total tau concentrations, which are markers of neuronal and axonal cell injury and neurofibrillary tangles (Suarez-Calvet *et al*., 2019). We show that the elevation of CSF sTREM2 levels could differentiate PD participants with or without abnormal CSF tau concentration with an area under the curve of 0.828 (CI = 0.757-0.899).

These results also suggest that CSF sTREM2 may serve as a general biomarker of neuronal injury across neurological diseases characterized by a neuroinflammatory component. Beyond PD and AD, CSF sTREM2 is also elevated in HIV-1 infection (Gisslen *et al*., 2019), where high levels of hyperphosphorylated tau protein have been documented (Brew *et al*., 2005). CSF sTREM2 concentration also is elevated in relapsing-remitting multiple sclerosis (MS), primary progressive MS, and other inflammatory neurological disease (OIND) subjects (Piccio *et al*., 2008), and these forms of MS are also associated with tau pathology (Bartosik-Psujek and Stelmasiak, 2006; Anderson *et al*., 2008; Anderson *et al*., 2010; Jaworski *et al*., 2012). We also found a positive association between CSF sTREM2 and Aβ_1-40_ concentrations across all diagnostic groups, an association previously observed in dementia patients (Henjum *et al*., 2018).

sTREM2 is produced either from cleavage of surface-expressed TREM2 on brain microglia and parenchymal macrophages, or from alternative splicing in those cells; the relative proportion of these contributions is unknown, as it is not possible to estimate levels of surface TREM2 at this time in brain. If sTREM2 is generated mainly through cleavage of functional surface TREM2, that would indicate that the beneficial TREM2 immune response is in decline, heralding progression to full dementia. Indeed, the findings in this study and those in AD, where sTREM2 peaks at the MCI stage and then declines, may be consistent with the idea that compensatory anti-inflammatory and pro-phagocytic TREM2 responses become limiting, allowing disease-causing processes to accelerate.

In plasma, we observed a 2.1-fold increase of sTREM2 concentration compared to CSF, however, no significant difference was observed in any of the PD subgroups. This is possibly due to the cross-sectional nature of this study and the high level of variance in plasma sTREM2. Follow-up studies with longitudinal sampling may reduce the variance observed. For example, in a prospective longitudinal study, higher levels of sTREM2 in blood were associated with increased risk of developing dementia, AD, and vascular dementia in the general elderly Japanese population (Ohara *et al*., 2019).

Finally, the finding that CSF sTREM2 is positively associated with MoCA score in PD participants with elevated CSF tau concentration is of interest. A putative neuroprotective association of CSF sTREM2 suggests an increase in functioning, surface expressed TREM2 in brain either before or at the time sTREM2 is elevated in CSF. A neuroprotective function of TREM2 signaling in microglia is supported by genome wide association studies (Guerreiro *et al*., 2013; Jonsson *et al*., 2013) and preclinical research in mouse models of AD (Wang *et al*., 2016; Raha *et al*., 2017; Ulland *et al*., 2017; Cheng-Hathaway *et al*., 2018; Parhizkar *et al*., 2019). A neuroprotective role of sTREM2 has been recently proposed in the 5xFAD mouse AD model where direct injection of sTREM2 protein or viral-mediated overexpression of sTREM2 in hippocampus was associated with a reduction of amyloid plaque load and a rescue of spatial memory and long-term potentiation deficits (Zhong *et al*., 2019). It is possible that increasing CSF sTREM2 levels in advancing cognitive decline might reflect prior or ongoing TREM2 signaling that is being terminated by cleavage of the membrane bound TREM2 and release. Further studies are warranted to better understand the temporal relationship between parenchymal microglial TREM2 expression and signaling and the appearance of the soluble form of TREM2.

In conclusion, we demonstrate that levels of CSF sTREM2 increase in PD patient subgroups with abnormal CSF tau concentration, with or without abnormal CSF amyloid concentration, and this increase is observed in cognitively normal and MCI PD subgroups. These findings suggest that CSF sTREM2 could serve as a surrogate biomarker of TREM2-mediated microglia function and neuronal injury in PD.

## References

Aarsland D, Andersen K, Larsen JP, Lolk A, Kragh-Sorensen P. Prevalence and characteristics of dementia in Parkinson disease: an 8-year prospective study. Arch Neurol 2003; 60(3): 387–92.

Aarsland D, Bronnick K, Larsen JP, Tysnes OB, Alves G, Norwegian ParkWest Study G. Cognitive impairment in incident, untreated Parkinson disease: the Norwegian ParkWest study. Neurology 2009; 72(13): 1121–6.

Anderson JM, Hampton DW, Patani R, Pryce G, Crowther RA, Reynolds R, et al. Abnormally phosphorylated tau is associated with neuronal and axonal loss in experimental autoimmune encephalomyelitis and multiple sclerosis. Brain 2008; 131(Pt 7): 1736–48.

Anderson JM, Patani R, Reynolds R, Nicholas R, Compston A, Spillantini MG, et al. Abnormal tau phosphorylation in primary progressive multiple sclerosis. Acta Neuropathol 2010; 119(5): 591–600.

Bartosik-Psujek H, Stelmasiak Z. The CSF levels of total-tau and phosphotau in patients with relapsing-remitting multiple sclerosis. J Neural Transm (Vienna) 2006; 113(3): 339–45.

Bayart JL, Hanseeuw B, Ivanoiu A, van Pesch V. Analytical and clinical performances of the automated Lumipulse cerebrospinal fluid Abeta42 and T-Tau assays for Alzheimer’s disease diagnosis. J Neurol 2019.

Benitez BA, Cooper B, Pastor P, Jin SC, Lorenzo E, Cervantes S, et al. TREM2 is associated with the risk of Alzheimer’s disease in Spanish population. Neurobiol Aging 2013; 34(6): 1711 e15–7.

Brew BJ, Pemberton L, Blennow K, Wallin A, Hagberg L. CSF amyloid beta42 and tau levels correlate with AIDS dementia complex. Neurology 2005; 65(9): 1490–2.

Buchman AS, Yu L, Wilson RS, Leurgans SE, Nag S, Shulman JM, et al. Progressive parkinsonism in older adults is related to the burden of mixed brain pathologies. Neurology 2019; 92(16): e1821–e30.

Chen-Plotkin AS, Albin R, Alcalay R, Babcock D, Bajaj V, Bowman D, et al. Finding useful biomarkers for Parkinson’s disease. Sci Transl Med 2018; 10(454).

Cheng-Hathaway PJ, Reed-Geaghan EG, Jay TR, Casali BT, Bemiller SM, Puntambekar SS, et al. The Trem2 R47H variant confers loss-of-function-like phenotypes in Alzheimer’s disease. Mol Neurodegener 2018; 13(1): 29.

Cholerton B, Johnson CO, Fish B, Quinn JF, Chung KA, Peterson-Hiller AL, et al. Sex differences in progression to mild cognitive impairment and dementia in Parkinson’s disease. Parkinsonism Relat Disord 2018; 50: 29–36.

Cholerton BA, Zabetian CP, Quinn JF, Chung KA, Peterson A, Espay AJ, et al. Pacific Northwest Udall Center of excellence clinical consortium: study design and baseline cohort characteristics. J Parkinsons Dis 2013; 3(2): 205–14.

Chou KL, Amick MM, Brandt J, Camicioli R, Frei K, Gitelman D, et al. A recommended scale for cognitive screening in clinical trials of Parkinson’s disease. Movement disorders: official journal of the Movement Disorder Society 2010; 25(15): 2501–7.

Colonna M. TREMs in the immune system and beyond. Nature reviews Immunology 2003; 3(6): 445–53.

de Lau LM, Breteler MM. Epidemiology of Parkinson’s disease. Lancet Neurol 2006; 5(6): 525–35.

Dickson DW, Heckman MG, Murray ME, Soto AI, Walton RL, Diehl NN, et al. APOE epsilon4 is associated with severity of Lewy body pathology independent of Alzheimer pathology. Neurology 2018; 91(12): e1182–e95.

Emre M, Mecocci P, Stender K. Pooled analyses on cognitive effects of memantine in patients with moderate to severe Alzheimer’s disease. J Alzheimers Dis 2008; 14(2): 193–9.

Gisslen M, Heslegrave A, Veleva E, Yilmaz A, Andersson LM, Hagberg L, et al. CSF concentrations of soluble TREM2 as a marker of microglial activation in HIV-1 infection. Neurol Neuroimmunol Neuroinflamm 2019; 6(1): e512.

Guerreiro R, Wojtas A, Bras J, Carrasquillo M, Rogaeva E, Majounie E, et al. TREM2 variants in Alzheimer’s disease. The New England journal of medicine 2013; 368(2): 117–27.

Hamerman JA, Jarjoura JR, Humphrey MB, Nakamura MC, Seaman WE, Lanier LL. Cutting edge: inhibition of TLR and FcR responses in macrophages by triggering receptor expressed on myeloid cells (TREM)-2 and DAP12. J Immunol 2006; 177(4): 2051–5.

Hely MA, Reid WG, Adena MA, Halliday GM, Morris JG. The Sydney multicenter study of Parkinson’s disease: the inevitability of dementia at 20 years. Mov Disord 2008; 23(6): 837–44.

Hendershott TR, Zhu D, Llanes S, Poston KL. Domain-specific accuracy of the Montreal Cognitive Assessment subsections in Parkinson’s disease. Parkinsonism Relat Disord 2017; 38: 31–4.

Hendershott TR, Zhu D, Llanes S, Zabetian CP, Quinn J, Edwards KL, et al. Comparative sensitivity of the MoCA and Mattis Dementia Rating Scale-2 in Parkinson’s disease. Mov Disord 2019; 34(2): 285–91.

Henjum K, Almdahl IS, Arskog V, Minthon L, Hansson O, Fladby T, et al. Cerebrospinal fluid soluble TREM2 in aging and Alzheimer’s disease. Alzheimer’s research & therapy 2016; 8(1): 17.

Henjum K, Quist-Paulsen E, Zetterberg H, Blennow K, Nilsson LNG, Watne LO. CSF sTREM2 in delirium-relation to Alzheimer’s disease CSF biomarkers Abeta42, t-tau and p-tau. J Neuroinflammation 2018; 15(1): 304.

Heslegrave A, Heywood W, Paterson R, Magdalinou N, Svensson J, Johansson P, et al. Increased cerebrospinal fluid soluble TREM2 concentration in Alzheimer’s disease. Molecular neurodegeneration 2016; 11: 3.

Jack CR, Jr., Bennett DA, Blennow K, Carrillo MC, Dunn B, Haeberlein SB, et al. NIA-AA Research Framework: Toward a biological definition of Alzheimer’s disease. Alzheimers Dement 2018; 14(4): 535–62.

Jaworski J, Psujek M, Janczarek M, Szczerbo-Trojanowska M, Bartosik-Psujek H. Total-tau in cerebrospinal fluid of patients with multiple sclerosis decreases in secondary progressive stage of disease and reflects degree of brain atrophy. Ups J Med Sci 2012; 117(3): 284–92.

Jonsson T, Stefansson H, Steinberg S, Jonsdottir I, Jonsson PV, Snaedal J, et al. Variant of TREM2 associated with the risk of Alzheimer’s disease. The New England journal of medicine 2013; 368(2): 107–16.

Kleinberger G, Yamanishi Y, Suarez-Calvet M, Czirr E, Lohmann E, Cuyvers E, et al. TREM2 mutations implicated in neurodegeneration impair cell surface transport and phagocytosis. Science translational medicine 2014; 6(243): 243ra86.

Lewczuk P, Riederer P, O’Bryant SE, Verbeek MM, Dubois B, Visser PJ, et al. Cerebrospinal fluid and blood biomarkers for neurodegenerative dementias: An update of the Consensus of the Task Force on Biological Markers in Psychiatry of the World Federation of Societies of Biological Psychiatry. World J Biol Psychiatry 2018; 19(4): 244–328.

Marek K, Chowdhury S, Siderowf A, Lasch S, Coffey CS, Caspell-Garcia C, et al. The Parkinson’s progression markers initiative (PPMI) - establishing a PD biomarker cohort. Ann Clin Transl Neurol 2018; 5(12): 1460–77.

Muslimovic D, Post B, Speelman JD, Schmand B. Cognitive profile of patients with newly diagnosed Parkinson disease. Neurology 2005; 65(8): 1239–45.

Nasreddine ZS, Phillips NA, Bedirian V, Charbonneau S, Whitehead V, Collin I, et al. The Montreal Cognitive Assessment, MoCA: a brief screening tool for mild cognitive impairment. J Am Geriatr Soc 2005; 53(4): 695–9.

Neumann H, Takahashi K. Essential role of the microglial triggering receptor expressed on myeloid cells-2 (TREM2) for central nervous tissue immune homeostasis. J Neuroimmunol 2007; 184(1-2): 92–9.

Ohara T, Hata J, Tanaka M, Honda T, Yamakage H, Yoshida D, et al. Serum Soluble Triggering Receptor Expressed on Myeloid Cells 2 as a Biomarker for Incident Dementia: The Hisayama Study. Ann Neurol 2019; 85(1): 47–58.

Otero K, Turnbull IR, Poliani PL, Vermi W, Cerutti E, Aoshi T, et al. Macrophage colony-stimulating factor induces the proliferation and survival of macrophages via a pathway involving DAP12 and beta-catenin. Nat Immunol 2009; 10(7): 734–43.

Paciotti S, Sepe FN, Eusebi P, Farotti L, Cataldi S, Gatticchi L, et al. Diagnostic performance of a fully automated chemiluminescent enzyme immunoassay for Alzheimer’s disease diagnosis. Clin Chim Acta 2019; 494: 74–8.

Parhizkar S, Arzberger T, Brendel M, Kleinberger G, Deussing M, Focke C, et al. Loss of TREM2 function increases amyloid seeding but reduces plaque-associated ApoE. Nat Neurosci 2019; 22(2): 191–204.

Piccio L, Buonsanti C, Cella M, Tassi I, Schmidt RE, Fenoglio C, et al. Identification of soluble TREM-2 in the cerebrospinal fluid and its association with multiple sclerosis and CNS inflammation. Brain: a journal of neurology 2008; 131(Pt 11): 3081–91.

Piccio L, Deming Y, Del-Aguila JL, Ghezzi L, Holtzman DM, Fagan AM, et al. Cerebrospinal fluid soluble TREM2 is higher in Alzheimer disease and associated with mutation status. Acta Neuropathol (Berl) 2016.

Raha AA, Henderson JW, Stott SR, Vuono R, Foscarin S, Friedland RP, et al. Neuroprotective Effect of TREM-2 in Aging and Alzheimer’s Disease Model. J Alzheimers Dis 2017; 55(1): 199–217.

Rayaprolu S, Mullen B, Baker M, Lynch T, Finger E, Seeley WW, et al. TREM2 in neurodegeneration: evidence for association of the p.R47H variant with frontotemporal dementia and Parkinson’s disease. Mol Neurodegener 2013; 8: 19.

Ren M, Guo Y, Wei X, Yan S, Qin Y, Zhang X, et al. TREM2 overexpression attenuates neuroinflammation and protects dopaminergic neurons in experimental models of Parkinson’s disease. Exp Neurol 2018; 302: 205–13.

Robinson JL, Lee EB, Xie SX, Rennert L, Suh E, Bredenberg C, et al. Neurodegenerative disease concomitant proteinopathies are prevalent, age-related and APOE4-associated. Brain 2018; 141(7): 2181–93.

Suarez-Calvet M, Araque Caballero MA, Kleinberger G, Bateman RJ, Fagan AM, Morris JC, et al. Early changes in CSF sTREM2 in dominantly inherited Alzheimer’s disease occur after amyloid deposition and neuronal injury. Sci Transl Med 2016a; 8(369): 369ra178.

Suarez-Calvet M, Kleinberger G, Araque Caballero MA, Brendel M, Rominger A, Alcolea D, et al. sTREM2 cerebrospinal fluid levels are a potential biomarker for microglia activity in early-stage Alzheimer’s disease and associate with neuronal injury markers. EMBO Molecular Medicine 2016b; 8(5): 466–76.

Suarez-Calvet M, Morenas-Rodriguez E, Kleinberger G, Schlepckow K, Araque Caballero MA, Franzmeier N, et al. Early increase of CSF sTREM2 in Alzheimer’s disease is associated with tau related-neurodegeneration but not with amyloid-beta pathology. Mol Neurodegener 2019; 14(1): 1.

Tsuang D, Leverenz JB, Lopez OL, Hamilton RL, Bennett DA, Schneider JA, et al. APOE epsilon4 increases risk for dementia in pure synucleinopathies. JAMA Neurol 2013; 70(2): 223–8.

Turnbull IR, Gilfillan S, Cella M, Aoshi T, Miller M, Piccio L, et al. Cutting edge: TREM-2 attenuates macrophage activation. J Immunol 2006; 177(6): 3520–4.

Ulland TK, Song WM, Huang SC, Ulrich JD, Sergushichev A, Beatty WL, et al. TREM2 Maintains Microglial Metabolic Fitness in Alzheimer’s Disease. Cell 2017; 170(4): 649–63 e13.

van Steenoven I, Aarsland D, Weintraub D, Londos E, Blanc F, van der Flier WM, et al. Cerebrospinal Fluid Alzheimer’s Disease Biomarkers Across the Spectrum of Lewy Body Diseases: Results from a Large Multicenter Cohort. J Alzheimers Dis 2016; 54(1): 287–95.

Wang Y, Ulland TK, Ulrich JD, Song W, Tzaferis JA, Hole JT, et al. TREM2-mediated early microglial response limits diffusion and toxicity of amyloid plaques. The Journal of Experimental Medicine 2016.

Wunderlich P, Glebov K, Kemmerling N, Tien NT, Neumann H, Walter J. Sequential proteolytic processing of the triggering receptor expressed on myeloid cells-2 (TREM2) protein by ectodomain shedding and gamma-secretase-dependent intramembranous cleavage. The Journal of biological chemistry 2013; 288(46): 33027–36.

Yeh FL, Wang Y, Tom I, Gonzalez LC, Sheng M. TREM2 Binds to Apolipoproteins, Including APOE and CLU/APOJ, and Thereby Facilitates Uptake of Amyloid-Beta by Microglia. Neuron 2016; 91(2): 328–40.

Zhong L, Xu Y, Zhuo R, Wang T, Wang K, Huang R, et al. Soluble TREM2 ameliorates pathological phenotypes by modulating microglial functions in an Alzheimer’s disease model. Nat Commun 2019; 10(1): 1365.

